# Blocking Orai1 constitutive activity inhibits B-cell cancer migration and synergistically acts with drugs to reduce B-CLL cell survival

**DOI:** 10.1101/2023.12.12.571243

**Authors:** Julien Scaviner, Cristina Bagacean, Berthou Christian, Yves Renaudineau, Olivier Mignen, Souleymane Abdoul-Azize

**Affiliations:** INSERM UMR1227, Université de Bretagne Occidentale, F-29200 Brest, France

**Keywords:** B-cell cancers, pharmacological Orai1 blockers, Ca^2+^ signaling, migration, drug sensitivity

## Abstract

Orai1 channel capacity to control store-operated Ca^2+^ entry (SOCE) and B-cell functions is poorly understood and more specifically in B-cell cancers, including human lymphoma and leukemia. As compared to normal B-cells, Orai1 is overexpressed in B-chronic lymphocytic leukemia (B-CLL) and contributes in resting B-CLL to mediate an elevated basal Ca^2+^ level through a constitutive Ca^2+^ entry, and in BCR-activated B-cell to regulate the Ca^2+^ signaling response. Such observations were confirmed in human B-cell lymphoma and leukemia lines, including RAMOS, JOK-1, MEC-1 and JVM-3 cells. Next, the use of pharmacological Orai1 inhibitors (GSK-7975A and Synta66) blocks constitutive Ca^2+^ entry and in turn affects B-cell cancer (primary and cell lines) survival and migration, controls cell cycle, and induces apoptosis through a mitochondrial and caspase-3 independent pathway. Finally, the added value of Orai1 inhibitors in combination with B-CLL drugs (ibrutinib, idelalisib, rituximab, and venetoclax) on B-CLL survival was tested, showing an additive/synergistic effect including in the B-cell cancer lines. To conclude, this study highlights the pathophysiological role of the Ca^2+^ channel Orai1 in B-cell cancers, and pave the way for the use of ORAI1 modulators as a plausible therapeutic strategy.

## 1. Introduction

In response to an antigenic stimulation, lymphocyte cell signaling and activation are primarily governed by an increase of the cytoplasmic calcium (Ca^2+^) level [1]. Indeed, following B cell receptor (BCR) or T cell receptor (TCR) engagement, the increase of cytoplasmic Ca^2+^ depends first from the release of Ca^2+^ from the endoplasmic reticulum (ER) stores (Ca^2+^ channel independent process) followed by a concomitant influx of Ca^2+^ from the extracellular space [2,3]. In T and B cells,, Ca^2+^ entry is mainly driven by the activity of plasma membrane CRAC channels, which constitute the store-operated calcium entry (SOCE) [1,4]. CRAC channel-mediated Ca^2+^ entry regulates many immune cell functions by activating or inhibiting various signaling pathways through Ca^2+^-dependent proteins or transcription factors, including NFAT, nuclear factor κB (NF -κB) [5–7]. In addition to the SOCE, a constitutive and basal Ca^2+^ pathway has been reported in tumoral lymphocytes allowing their survival [8].

Among the three isoforms of Orai (Orai1, 2, 3), Orai1, also known as CRACM1, has long been described as the central component of CRAC channels [9]. Mutation causing a loss of function of Orai1 is associated with CRAC channelopathy that abolishes SOCE in T cells with symptoms combining immunodeficiency, muscle hypotonia, ectodermal dysplasia, and autoimmunity [10–12]. Unlike T cells, the role of Orai1 in B cells is much less known and results from Orai1^-/-^ knockout mice model supported that SOCE and in turn cell proliferation are significantly attenuated in response to BCR stimulation [13,14], suggesting that Orai1 contributes at least in part to Ca^2+^ signaling and effector functions in B cells. By contrast, the role of Orai1 in B cell leukemia remains unknown [15].

B-cell cancer such as chronic lymphocytic leukemia (CLL) is a cancer characterized by a dysregulated Ca^2+^ signaling pathway that governs the clinical outcomes of this pathology [16,17]. Although CLL could be considered to be a proliferative disease, the treatments used in B-CLL directly or indirectly affect Ca^2+^ signaling via BCR signaling (ibrutinib) [16], PI3K signaling (idelalisib) [18,19], Bcl-2 which interfere with the IP_3_-receptor (venetoclax) [20], or the CD20 receptor whose activation has been described as modulating Ca^2+^ signaling (rituximab) [21].

In this study, we investigated the Orai1 channel contribution in B-cell cancer from human primary specimens and derived cell lines to Ca^2+^ signaling, B cell functions, and the added value of Orai inhibitors to promote B-CLL-drug capacity to induce B-cell cancer death. Main reports from this study are related to the observations that: (i) Orai1 is overexpressed in primary B-CLL cells and malignant B-cell lines; (ii) Orai1 controls both the basal Ca^2+^ entry and the BCR dependent SOCE signaling; (iii) blocking Orai1 channel with pharmacological inhibitors affects B-cell cancer survival, mitochondrial functions and B-cell cancer line migration; and (iv) inhibition of Orai1 acts synergistically with CLL-drugs to reduce both primary specimens and B-cell cancer survival.

## 2. Results

### Orai1 is overexpressed in B-cell cancers and mediates Ca^2+^ signaling in primary B-CLL cells and B-cell cancer lines

Using murine primary B cells, it was recently demonstrated that Orai1 mediates CRAC function in B cells [14]. Accordingly, we wondered whether Orai1 might play the same role in human B-cell cancers and for that we purified B-cells from five CLL patients (Table1) and healthy controls, and selected four human B-cell cancers, including human B-CLL cell line MEC-1, B-cell prolymphocytic leukemia JVM-3, hairy B cell leukemia line JOK-1 and Burkitt lymphoma cell line RAMOS. Interestingly, Orai1 cell surface expression was investigated by flow cytometry showing an overexpression of Orai1 in the four B-cell cancer derived cell lines as compared to normal B cells (MFI Orai1: 0.69⍰±⍰0.30 in normal B cells versus 2.96 ±⍰1.50 in B-cell cancer lines, p=0.05) (Figure 1A, B), and an overexpression of Orai1 in primary B-CLL as compared to normal B cells (MFI Orai1: 0.69⍰±⍰0.30 in normal B cells versus 1.47⍰±⍰0.32 in B-CLL cells, p=0.01) (Figure 1B). Next, we analyzed RNA-seq dataset from our laboratory of human CLL samples (n = 4) available at GSE176141. Purified B cells from 4 untreated CLL patients were tested at two time points (diagnosis and 3 years after diagnosis). We observed an increase in Orai1 expression after three years from diagnosis (paired t-test, p=0.0481) (Figure 1C).

**Figure 1:**
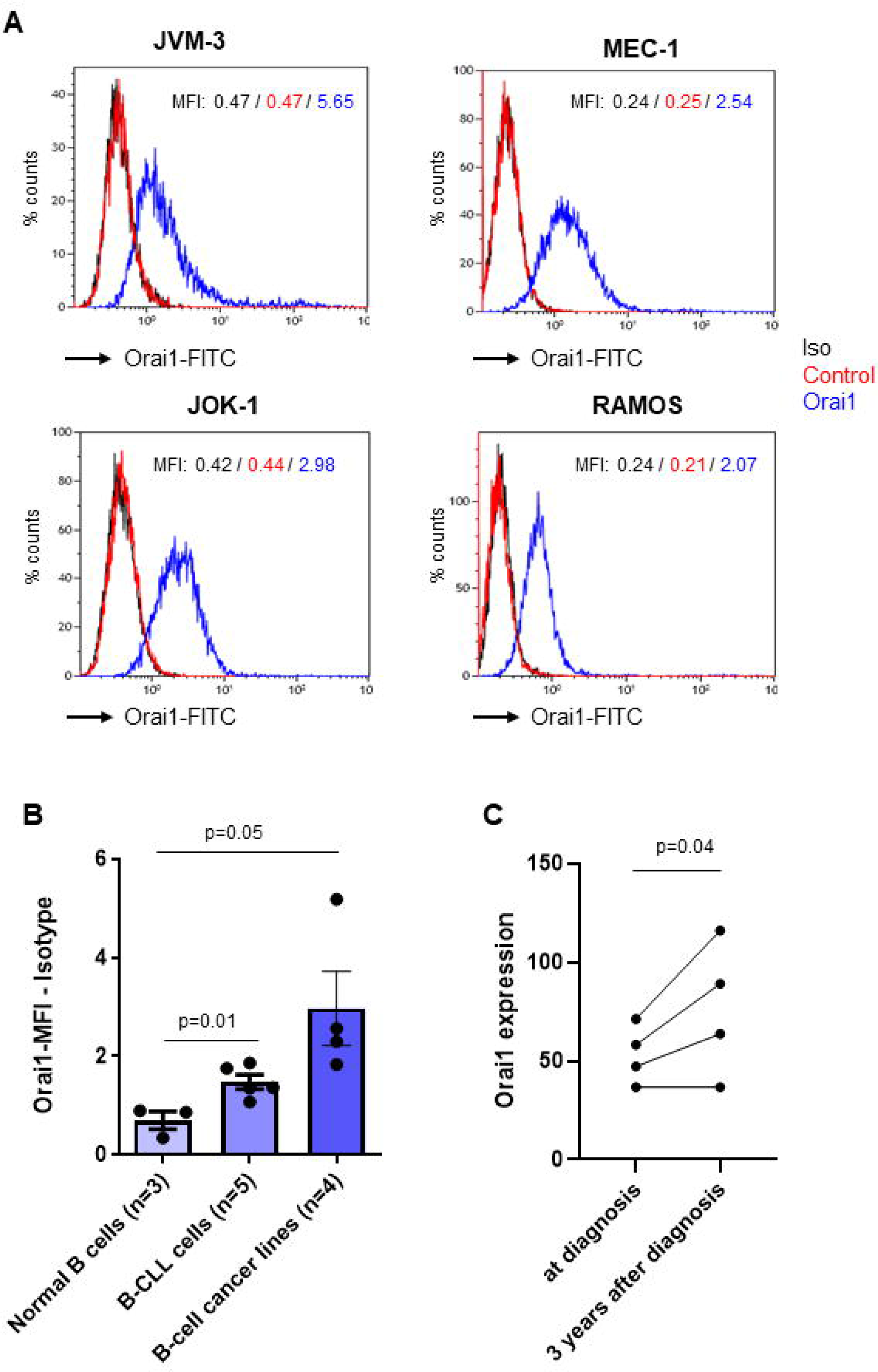
Orai1 expression in B-cell cancer lines and B-CLL primary cells. (A) Representative isotype (black), control (red) and Orai1 (blue) Flow cytometry histogram of B-cell cancer lines. Cells were stained with anti-Human Orai1 (extracellular)-FITC antibody. (B) Flow cytometry was performed to quantify Orai1 expression in mean fluorescence intensity (MFI, unpaired t-test) in normal B cells, B-CLL primary cells and B-cell cancer lines. (C) Differential expression of Orai1 in B-CLL primary cells isolated from patients at the time of disease diagnosis and three years after diagnosis with one-tailed paired t-test. Data were extracted from our RNA-seq data available at GSE176141.

Due to the importance of the Orai1 channel in the SOCE as previously demonstrated in primary mouse B cells [13], [14], [22], we investigated the contributions of Orai1 to Ca^2+^ mobilization following BCR engagement or activation. As shown in Figure 2, an increase in intracellular free Ca^2+^ concentrations ([Ca^2+^]_i_) is reported following BCR cross-link in the presence of anti-IgM Ab in B-cell cancer lines and such effect can be reversed in the presence of the two Orai1 channel blockers, namely Synta66 and GSK-7975A [23–25]. In B-CLL primary cells, anti-IgM Abs capacity to increase intracellular Ca^2+^ was similarly reduced in the presence of the two specific Orai1 inhibitor (Figure 2I, J). As one may argue that proximal BCR signaling is deficient in B-CLL from patients, and to confirm the role of orai1 in Ca^2+^ signaling of malignant B cells, the experiment was repeated in the presence of thapsigargin to by-pass the proximal BCR signal (TG, an inhibitor of the sarco/ER Ca^2+^-ATPase). Both inhibitors of Orai1 caused a near abrogation of SOCE in JOK-1, RAMOS and JVM-3 cell lines (Figure 3A-F) and a robust decrease in MEC-1 and primary cells of patients (Figure 3G-J). Figure 3K, L further confirmed Orai1 implication in SOCE B-cell cancer, indeed, when added after SOCE was initiated, blocking Orai1 with GSK-7975A significantly inhibited SOCE in JOK-1 cells.

**Figure 2:**
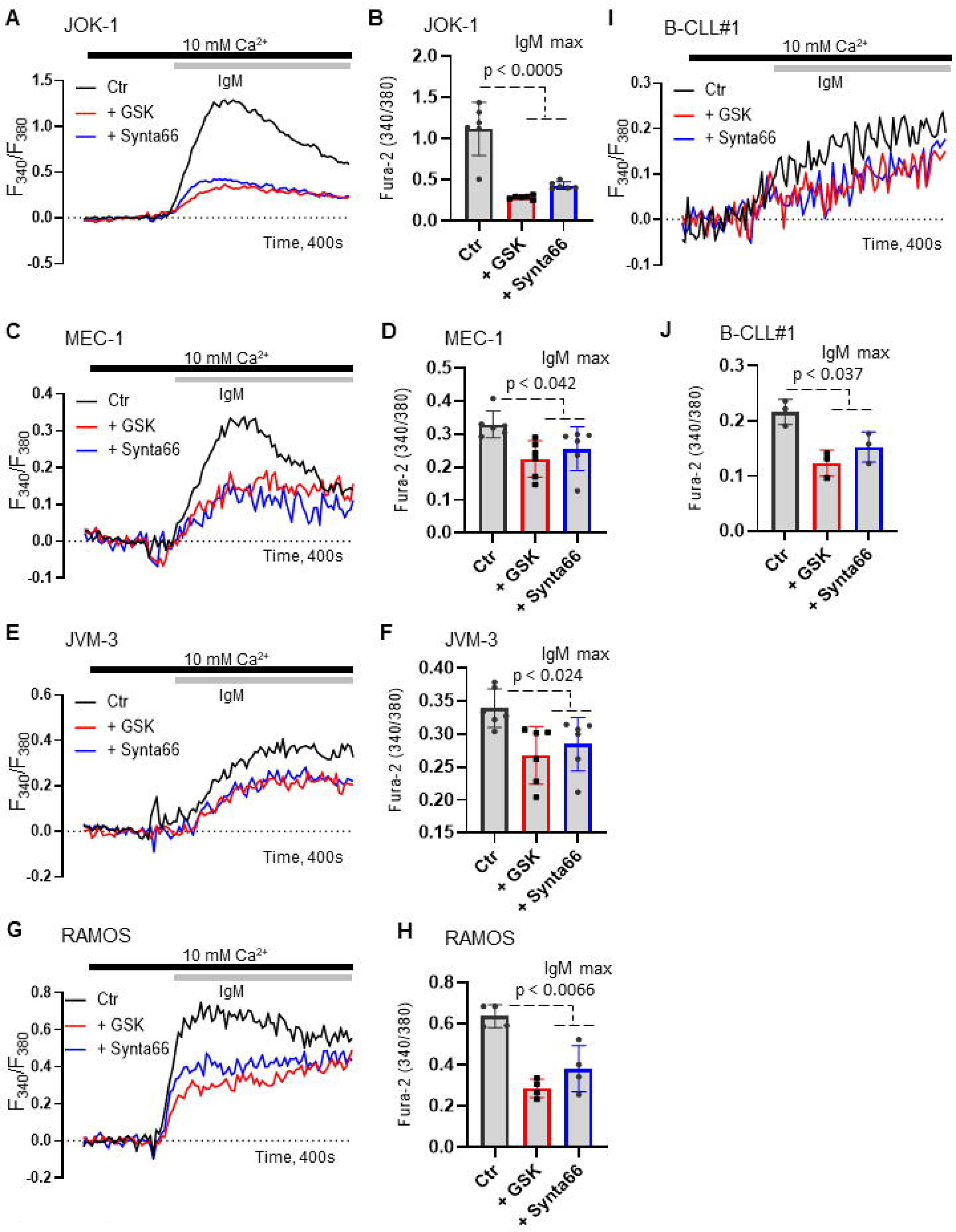
Orai1 inhibitors regulate BCR-mediated Ca^2+^ signaling in B-cell cancer derived cell lines. Cancer B-cells were preincubated during Fura-2 loading with GSK-7975A (GSK, 10 μM) or Synta66 (10 μM). (A, C, E, G, I) Representative kinetic plots of BCR-mediated Ca^2+^ signaling following IgM stimulation in 1.8 mM Ca^2+^. (B, D, F, H, J) Quantification of IgM peak in (A, C, E, G, I). All data in A-H are representative of at least 3 independent experiments. Statistical significance was analyzed by two-tailed, unpaired Student’s t test.

**Figure 3:**
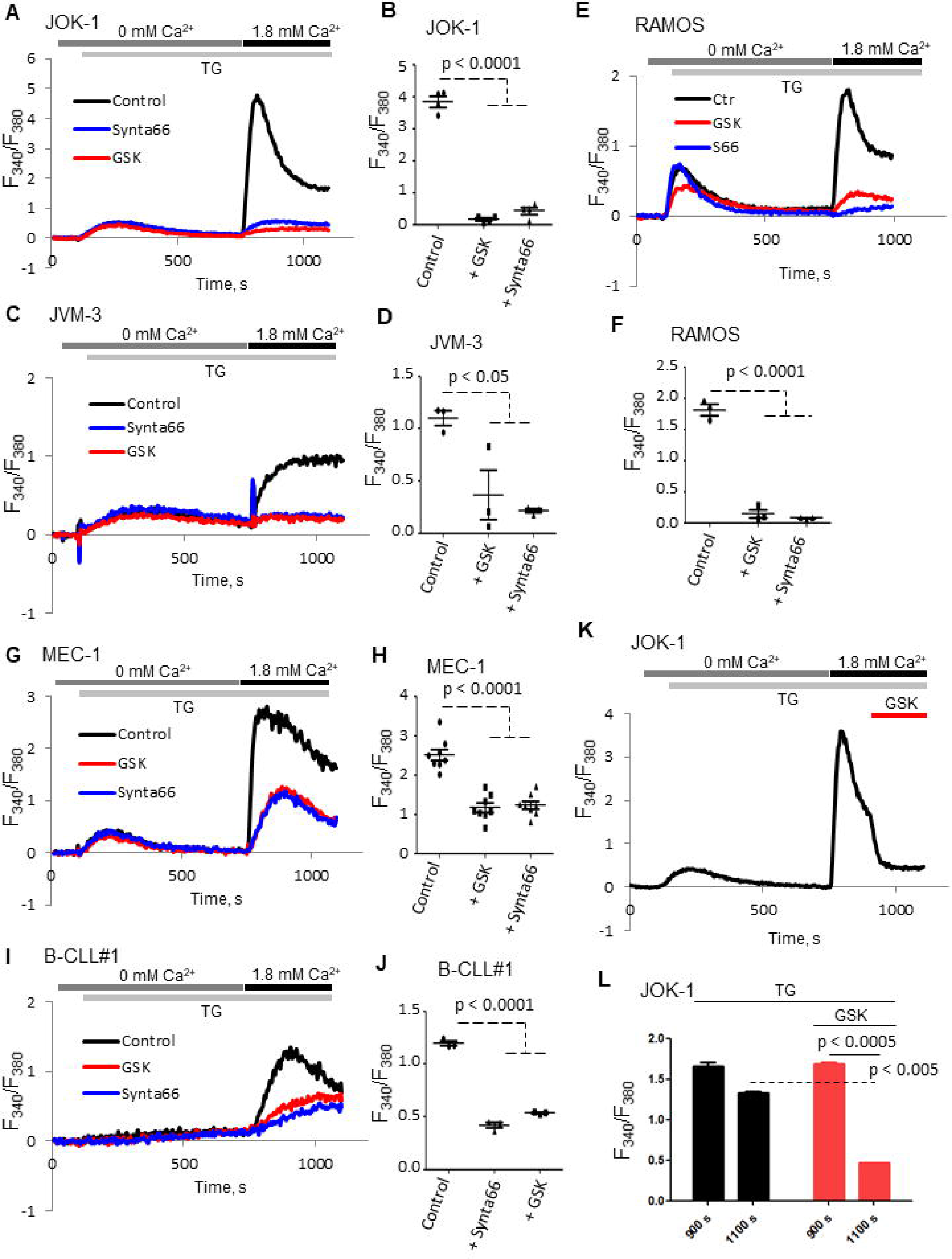
Orai1 inhibitors regulate store-operated Ca^2+^ entry (SOCE) in B-cell cancer lines and primary cells. (A, C, E, G, I) Representative traces of SOCE upon store depletion in 0 mM Ca^2+^ with 2 µM TG (thapsigargin) followed by addition of 1.8 mM Ca^2+^ to the solution. Cells before exposure to TG in Ca^2+^ buffer were preincubated during Fura-2 loading with GSK-7975A (GSK, 10 μM) or Synta66 (10 μM). (B, D, F, H, J) Quantification of SOCE peak in (A, C, E, G, I, K). (K) SOCE measurement as in (A) was inhibited with the addition of GSK-7975A (GSK, 10 μM) at 900 sec. (L) Quantification of peak SOCE in (K). With the exception of (I, J), all data are representative of at least 3 independent experiments. Statistical significance was analyzed by two-tailed, unpaired Student’s t test.

### Cancer B-cells display enhanced basal Orai1 channel activity

Ca^2+^ signaling deregulation in B-cell cancer progression was suggested [17] and based on our previous observation that a resting and constitutive Ca^2+^ entry pathway characterizes B-CLL from patients with progressive disease [26], we further hypothesize that such process may be related to Orai1 overexpression and activity. To this end, constitutive Ca^2+^ entry in B-cell cancer lines and B-CLL primary cells was evaluated by using the experimental method of Mn^2+^ quenching assay in the presence or not of Orai1 channel blockers. Results from such an analysis (Figure 4), reveals that both Orai1 channel blockers were sufficient to reduce constitutive Ca^2+^ entry (JOK: −21.75⍰±⍰0.57% reduction with GSK-7975A, −30.07±⍰8.31% with Synta66 compared to DMSO used as control; JVM-3: - 26.46⍰±⍰6.20% reduction with GSK-7975A, −60.36 ±⍰5.95% with Synta66 compared to control; MEC-1: −20.68⍰±⍰7.70% reduction with GSK-7975A, −26.01 ±⍰5.72% with Synta66 compared to control; RAMOS: −40.75⍰±⍰7.50% reduction with GSK-7975A, −48.35 ±⍰11.92% with Synta66 compared to control; Primary B-CLL : −44.33⍰±⍰7.33% reduction with GSK-7975A, −50.86 ±⍰2.73% with Synta66 compared to control).

**Figure 4:**
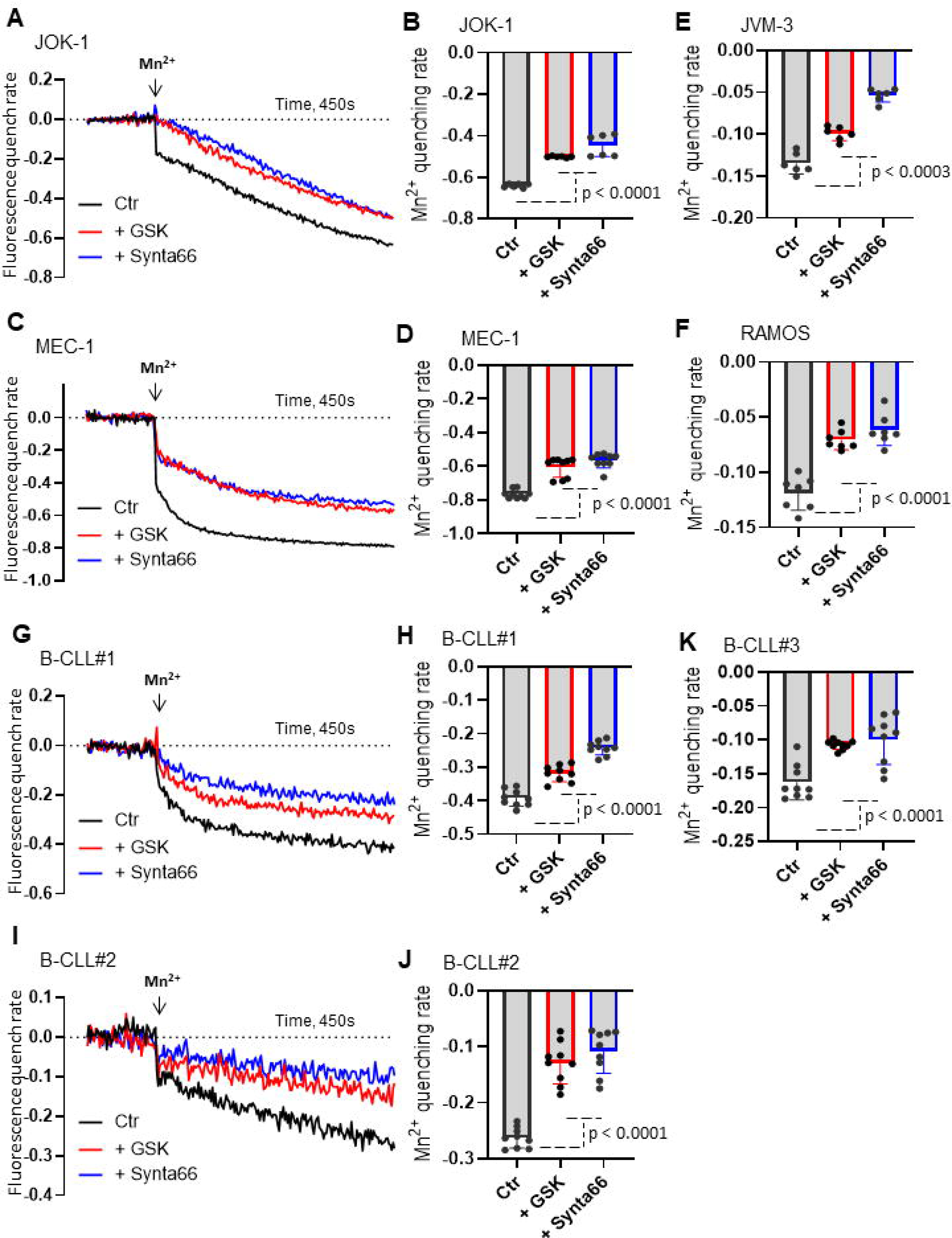
Orai1 blockers inhibit Mn^2+^ influx in B-cell cancer lines and primary cells. The quenching of Fura-2 fluorescence by Mn^2+^ in B-cell cancer lines was measured at 360 nm (the Ca^2+^ -independent excitation wavelength of Fura-2). (A, C, G, I) Cells before exposure to Mn^2+^ (10 μM) were preincubated during Fura-2 loading with GSK-7975A (GSK, 10 μM) or Synta66 (10 μM). (B, D, E, F, H, J, K) Quantification of quenching peak. Results express normalized fluorescence. Data are mean ± SEM (n > 3). Statistical significance was analyzed by two-tailed, unpaired Student’s t test.

Since this constitutive Ca^2+^ entry may impact the level of basal Ca^2+^ signaling described in B-CLL cells [27], we next characterize whether Orai1 activity is implicated in basal Ca^2+^ signaling in B-cell cancer and primary cells. To address this issue, basal Ca^2+^ level in five B-CLL primary samples and three normal human B cells were investigated. In agreement with constitutive Ca^2+^ entry results, B-CLL has high basal Ca^2+^ levels compared to normal B cells (Figure 5A), which was significantly reduced in the presence of Orai1 inhibitors compared to control (Ctr, DMSO) both in B CLL primary samples and B-cell cancer lines (Figure 5B-G). These results suggest that Orai1 contributes significantly to the constitutive Ca^2+^ entry and basal Ca^2+^ in resting B-cell cancers.

**Figure 5:**
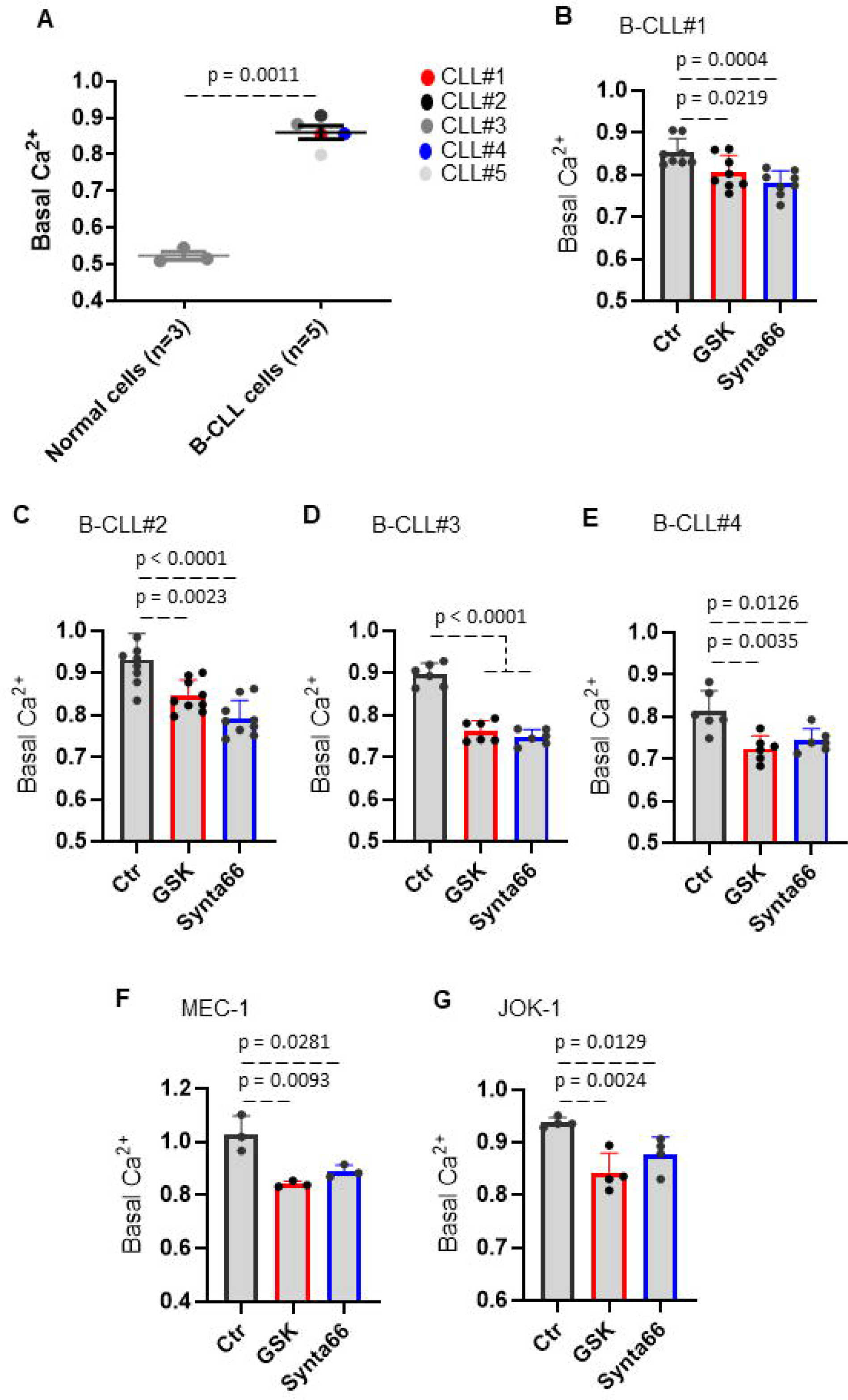
B-cell cancer lines and primary cells display constitutive Orai1 activity. The basal Ca^2+^ level was used as a read-out for measuring the constitutive Orai1 channel signaling in B-cell cancer. (A) Basal cytosolic Ca^2+^ level in primary normal B-cells compared to patient B-CLL cells. (B-G) Primary B-CLL cell and B-cell cancer lines pre-treated with control (Ctr, DMSO) and Orai1 blockers (GSK-7975A, GSK, 10 μM; or Synta66, 10 μM) using the ratiometric Ca^2+^ indicator Fura-2 AM. The basal Ca^2+^ level was evaluated before normalization as the average of initial F340nm/F380nm values. F, G data are representative of at least 3 independent experiments. Statistical significance was analyzed by two-tailed, unpaired Student’s t test.

### Blocking the Orai1 channel blocked B-cell cancer cell cycle and decreased cell proliferation

In contrast to B-CLL isolated from patients, B-cell cancer derived cell line have conserved their proliferative capacity and part of this activity may be related the overexpression of the Orai1 channel as Orai1 was reported to be critical for proliferation (when overexpressed) and apoptosis (when blocked) in mouse B cells [14,28]. We therefore, examined whether blocking Orai1 constitutive signal in the presence of GSK-7975A regulates cellular division in JOK-1 and MEC-1 B-cancer derived cells labeled with CellTrace Violet Fluorescence (CTV). The addition of GSK-7975A in the cell culture media was effective to reduce cell line proliferation in both JOK-1 (−63.89 ±⍰0.74% e.g. for GSK 50 µM) and MEC-1 (−13.89 ±⍰1.02% e.g. for GSK 50 µM) cell lines (Figure 6A, B). We also found that cell proliferation was significantly reduced by a 48-hours treatment with Synta66 in JOK-1 (−29.04 ±⍰1.78% e.g. for S66 20 µM) cell line (Figure 6C) as well as RAMOS cell line (−35.45 ±⍰10.39% e.g. for S66 20 µM) (Figure 6D). In order to understand the underlying regulatory mechanisms of this fundamental process, we investigated how Orai1 channel inhibition disturbs the cell cycle process in B-cell cancers. Cells were treated with Orai1 channel inhibitor, GSK-7975A, which promoted cell cycle arrest at the subG1 phase while negatively regulated the S phase (Figure 6E-J) in dose dependent manner, in accordance with our cell proliferation analysis determined by CTV and CCK-8 assays. Collectively, these findings demonstrate that Orai1 promotes proliferation signaling in B-cell cancer lines by regulating cell cycle progression and division.

**Figure 6:**
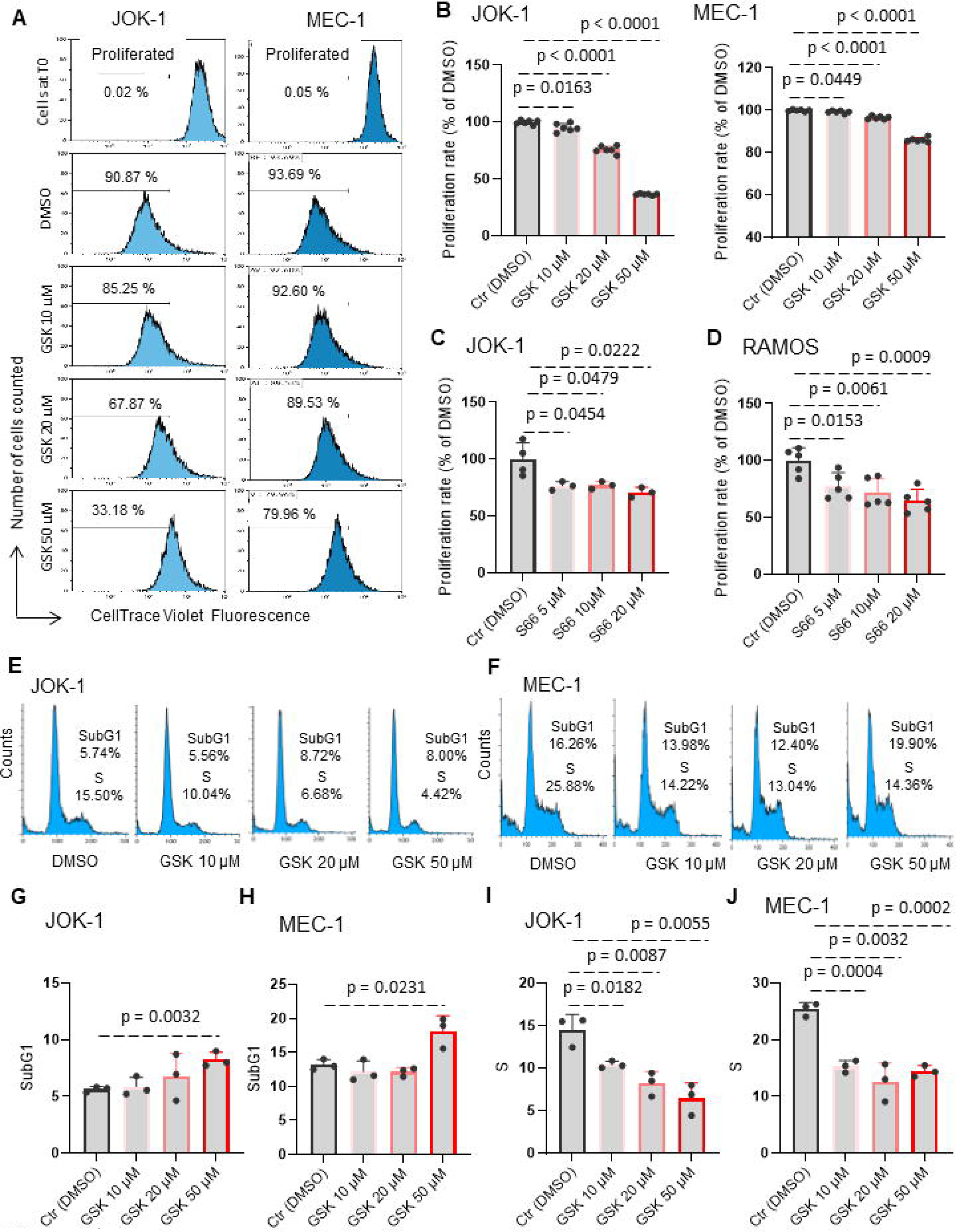
Orai1 inhibitor reduced proliferation by inhibiting the S phase of the cell cycle in B-cell cancer lines. (A, B) Measurement of B-cell cancer proliferation by tracking CellTrace Violet Fluorescence (CTV) dilution. Cells were loaded with CTV (1:1000 PBS) for 10 minutes and incubated with GSK-7975A (GSK, 10, 20, 50 μM). CTV dilution was determined 120 hours after. (A) Representative flow cytometry plots of B-cell cancer. (B) Quantification of the percentage of proliferating cells in (A). (C, D) Cells were treated with Synta66 (S66, 5, 10, 20 μM). Proliferation rate was determined by CCk-8 staining after 48 hours. The percentage was established after normalizing on control (Ctr) cells. (E-J) Measurement of B-CLL cell cycle phases. Cells were incubated with GSK-7975A (GSK, 10, 20, 50 μM). (E, F) Representative flow cytometry plots of JOK-1 and MEC-1 cell lines. (G-J) Quantification in (E, F) of the percentage of cell cycle phases, SubG1 phase (G, H) and S phase (I, J). This figure is representative of one of two independent experiments. Statistical significance was analyzed by two-tailed, unpaired Student’s t test.

### Orai1 inhibitors significantly increased B-cell cancer death in caspase-3 independent manner

After having established that the B-cell cancer cell cycle was regulated by the Orai1 signaling, and observed the increase in the number of SubG1 cells (Figure 6G,H), which are good indicators of apoptosis [29], three strategies were developed to better characterize the role of Orai1 channel in B-cell cancer death : i) Using LIVE/DEAD® viability dye, ii) annexin V-PI staining, iii) caspase-3 activity. First, treatment of B-CLL cells and B-cell cancer lines with Orai1 channel inhibitor, GSK-7975A, resulted in increased cell death, as manifested by a significant amount of dead cell population in red color (Figure 7A, C) and quantified in (Figure 7B, D). Second, the percentage of apoptotic cells, determined by annexin V-PI, was increased in the presence of Orai1 channel blocker (Figure 7E, F). In contrast, we observed no differences in caspase-3 activity in B-CLL cells and B-cell cancer lines (Figure 7G, H). These results show that blocking of Orai1 channel increased cell death in caspase-3-independent manner in B-cell malignancies.

**Figure 7:**
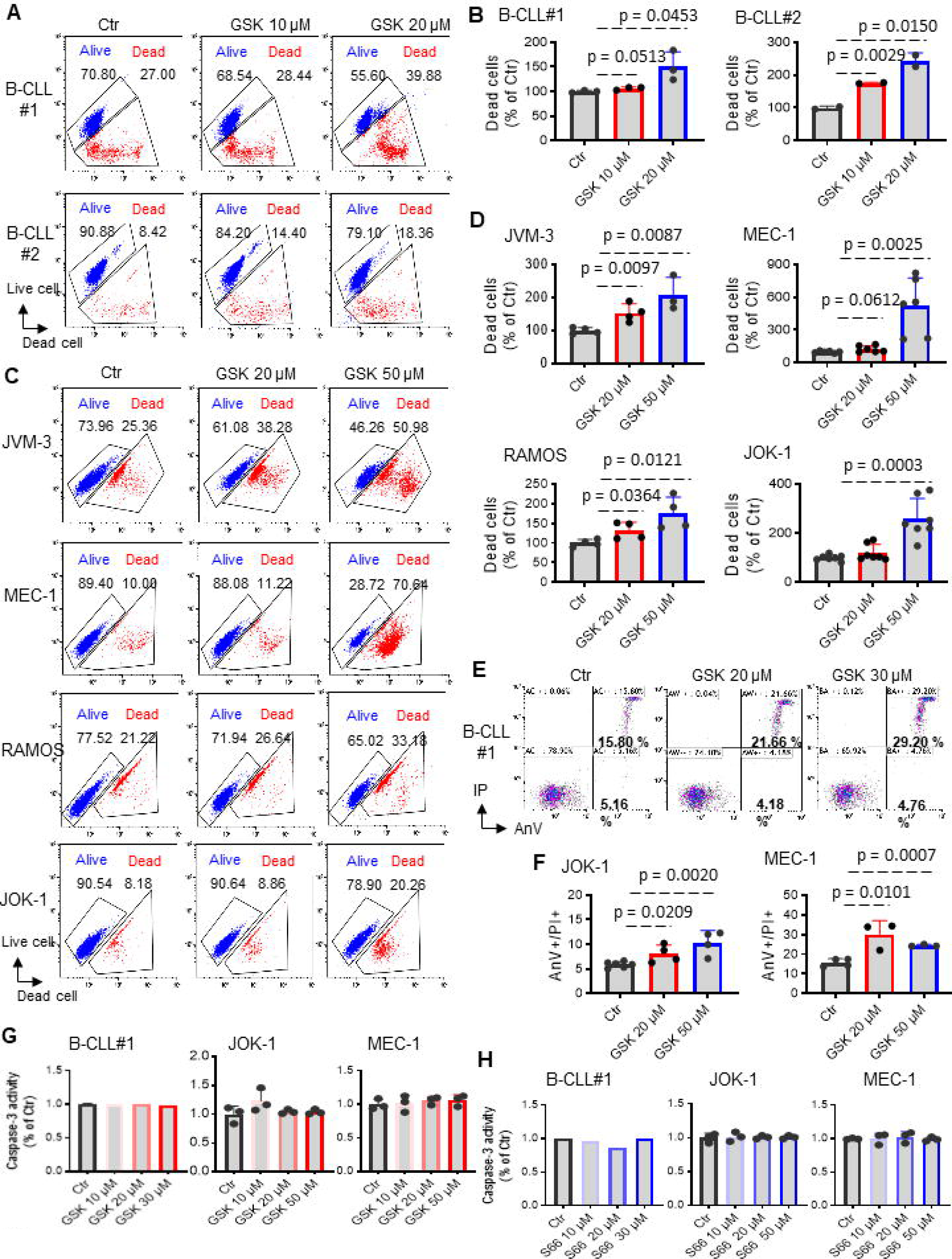
Orai1 inhibitor triggers cytotoxicity in B-cell cancer lines and primary cells. Primary B-CLL cells (A, B) and B-cell cancer lines (C, D) treated with GSK-7975A (GSK, 10, 20, 50 μM) for 48 hours. Alive and dead cells were determined using LIVE/DEAD® staining Kits and analyzed by flow cytometry. (A, C) Representative flow cytometry plots. (B, D) Quantification of the percentage of dead cells in (A, C). Cell mortality was also determined by flow cytometry after annexin V/PI staining (E, F). The percentage of total apoptotic cells was calculated as the percentage of annexin V-FITC positive plus annexin V-FITC/PI-positive population. (G, H) Caspase-3 activity determined using Ac-DEVD-AFC as substrate after 24 hours with GSK-7975A (GSK, 0-50 μM) or Synta66 (S66, 0-50 μM) in primary B-CLL cells and B-cell cancer lines. Data from B-cell cancer lines are representative of at least 3 independent experiments. Statistical significance was analyzed by two-tailed, unpaired Student’s t test.

### Orai1 inhibitors induced the loss of mitochondrial membrane potential in malignant B-cell lines

Because cell death can result from mitochondrial dysfunction [30], we next determined whether induction of cell death by blocking Orai1 channel was mediated through modulating mitochondrial functions. To this end, B-cell cancer lines and primary B-CLL cells were treated with or without Orai1 channel blockers for 24 hours. The mitochondrial membrane potential (Δψ_m_) dissipation, an early event for cell apoptosis, was detected by JC-10 (a lipophilic cationic dye) by flow cytometry. As shown in Figure 8A, B, we observed a significant increase in Δψm depolarization in cells treated with Orai1 inhibitors, in agreement with the cell death results.

**Figure 8:**
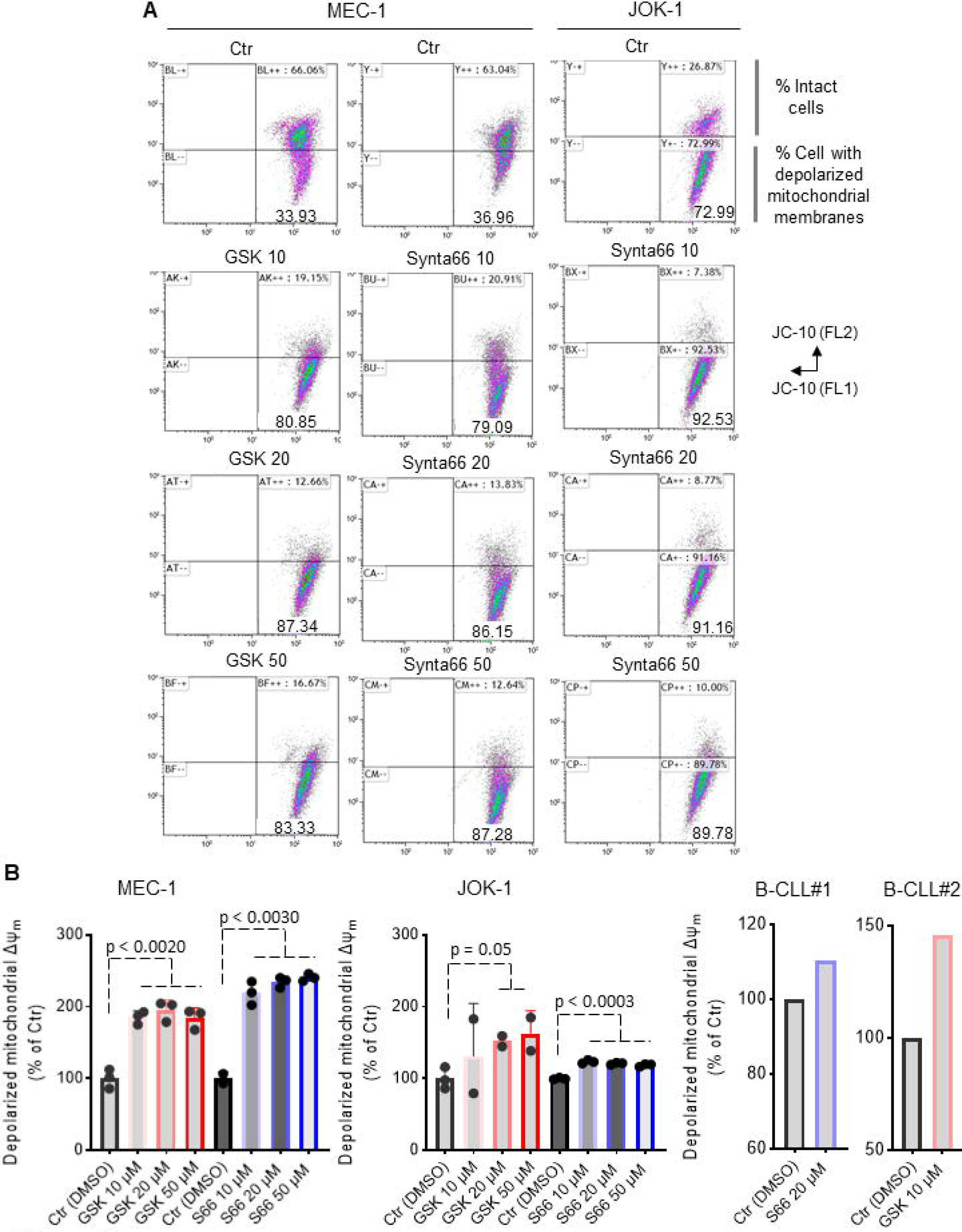
Orai1 inhibitor triggers the loss of mitochondrial membrane potential in B-cell cancer lines and primary cells. (A) Representative flow cytometry plots of B-cell cancer lines treated with GSK-7975A (GSK, 10, 20, 50 μM) or Synta66 (S66, 10, 20, 50 μM) for 24 hours showing mitochondria membrane potential change using JC-10 dye. (B) Quantification of cells with depolarized mitochondria from panel (A) and from primary B-CLL cells. Data are representative of at least 3 independent experiments in B-cell cancer lines. Statistical significance was analyzed by two-tailed, unpaired Student’s t test.

### Orai1 inhibitors reduced B-cell cancer migration in dose dependent manner

It is well known that Ca^2+^ signaling is a key regulator of tumor invasion [31]. Thus, several studies have shown the role of the Orai1 channel in the migration and metastasis of many tumor cells [32–37]. We next examined the effect of Orai1 channel bockers on SDF-1α-induced cell migration in MEC-1 and JOK-1 cell lines. We showed that GSK-7975A and Synta66 significantly inhibited B-leukemia derived cell line migration in response to SDF-1α, compared with control (DMSO) untreated cells and showed a dose-dependent effect (Figure 9A, B). Collectively, these results reveal that Orai1 channel contributes to B-cell cancer migration.

**Figure 9:**
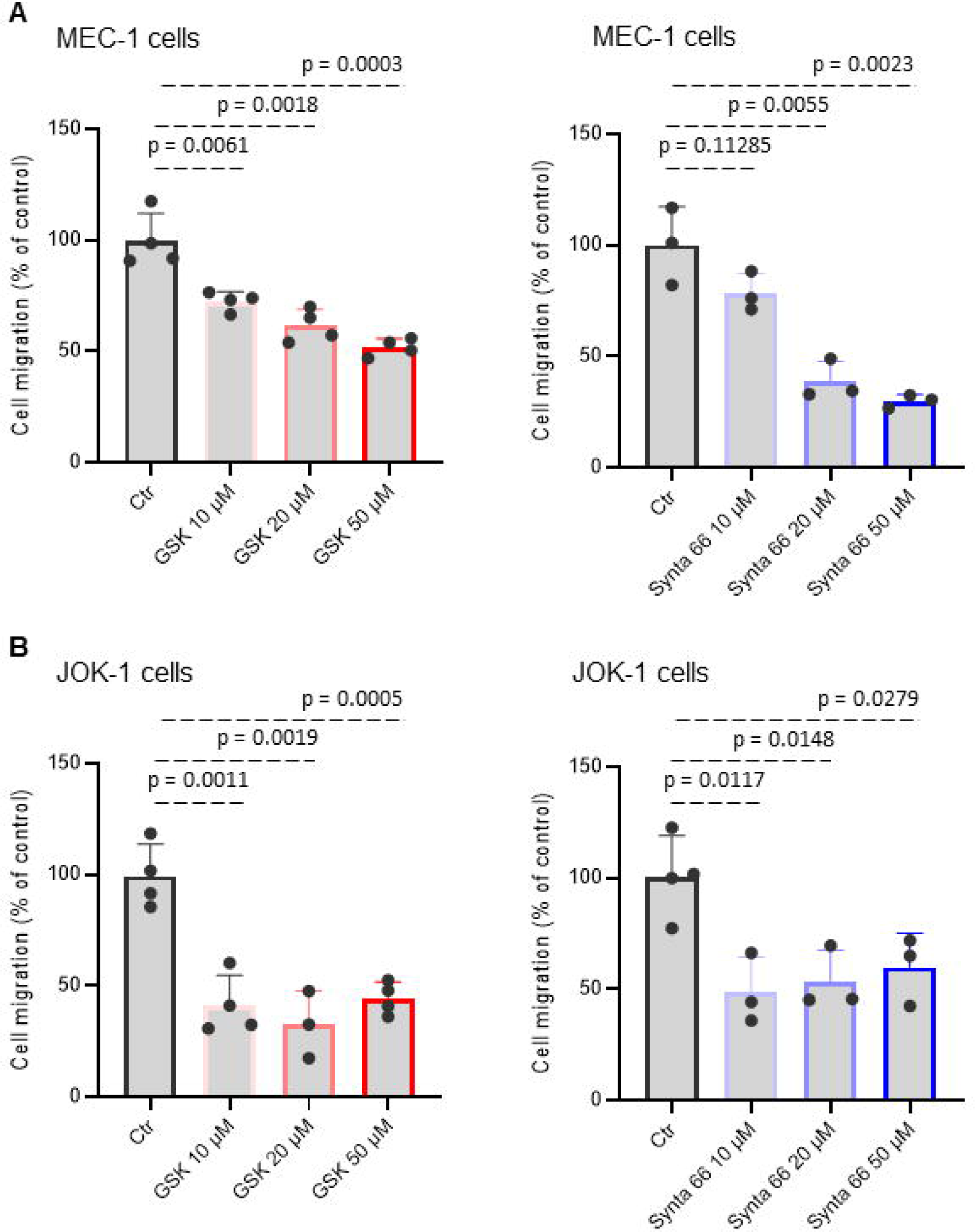
Orai1 inhibitors reduced B-cell leukemia line migration in dose dependent manner. Histograms showing B-cell leukemia migration treated with GSK-7975A (GSK, 10, 20, 50 μM) or Synta66 (10, 20, 50 μM) for 24 hours. The normalized cell number corresponds to the ratio of total number of migrating cells in presence of inhibitors/total number of migrating cells in control experiments. Data are representative of 2 independent experiments. Statistical significance was analyzed by two-tailed, unpaired Student’s t test.

### Orai1 inhibitors potentiated B-CLL drugs sensitivity in both B-CLL primary cells and B-cell cancer lines

Based on previous observation that blocking Orai1 channel was reported to improve rituximab-induced apoptosis in vitro in human non-hodgkin B lymphoma cells [38], we therefore evaluated the added value of Orai1 inhibitors in combination with B-CLL drugs (ibrutinib, idelalisib, rituximab, and venetoclax) on both B-CLL primary and B-cell cancer survival. We found that B-cell mortality determined by LIVE/DEAD® dye markedly increased by association of idelalisib (Idela) with inhibitors compared to Idela alone in four B-cell cancer lines as well as primary B-CLL cells (Figure 10A-E). These experiments were further repeated using annexin V-FITC/PI staining, showing that apoptosis rate was significantly increased by co-treatment in both B-leukemia cell lines JOK-1 (Figure 10F) and MEC-1 (Figure 10G), confirming the results obtained by the LIVE/DEAD® dye assay. Similar results showing an additive/synergistic effect including in the B-cancer cell lines were obtained by association of Orai1 inhibitors with ibrutinib (Ibrut) (Figure 10H-L), with rituximab (Rtx) (Figure 10M-Q) or with venetoclax (Vene) (Figure 10R-T). Interestingly, Orai1 blockers were sufficient to completely restore rituximab (Rtx) sensitivity in both resistant JOK-1 and MEC-1 cell lines (Figure 10M, N). These data suggest that Orai1 modulators could be a rational therapeutic strategy to improve the efficacy of chemotherapy in B-cell cancers.

**Figure 10:**
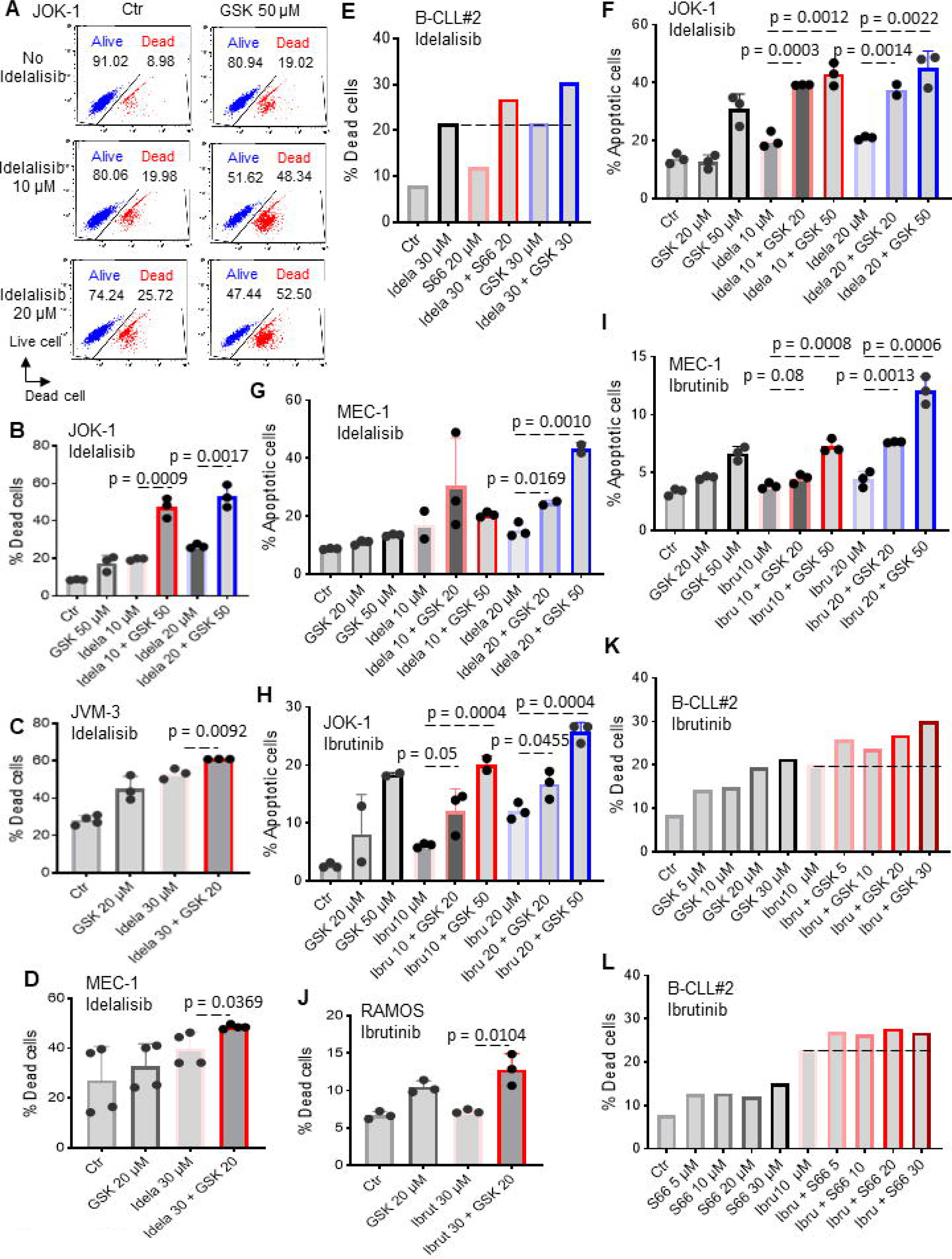
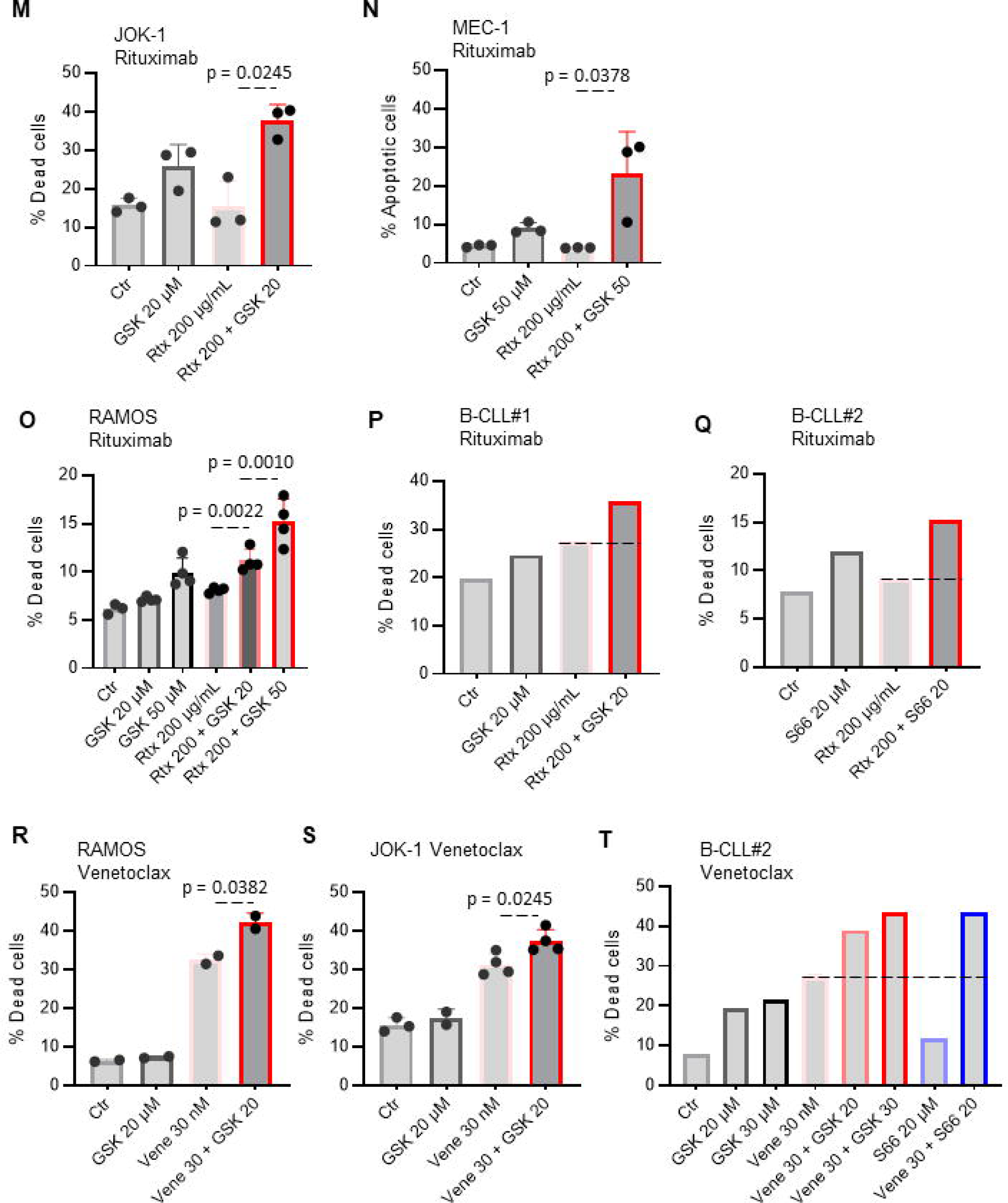
Orai1 inhibitors potentiated B-CLL drugs sensitivity in both primary and B-cell leukemia lines. Mortality of primary B-CLL cells and B-cell leukemia lines exposed for 48 hours to different drugs in the absence or presence of different concentrations of Orai1 inhibitors (GSK-7975A, GSK or Synta66, S66). (A) Representative flow cytometry plots showing alive and dead cells determined using LIVE/DEAD® dye. (B) Quantification of the percentage of dead cells in (A). Cell mortality following idelalisib treatment in JOK-1 cells (B), in JVM-3 cells (C), in MEC-1 cells (D), in primary B-CLL cells (E) determined by LIVE/DEAD® dye. Cell mortality was also determined by flow cytometry after annexin V/PI staining under Idelalisib (F, G) or ibrutinib (H, I) treatments. The percentage of total apoptotic cells was calculated as the percentage of annexin V-FITC positive plus annexin V-FITC/PI-positive population. Quantification of the percentage of dead cells following ibrutinib treatment in RAMOS cells (J), in primary B-CLL cells (K, L) determined by LIVE/DEAD® dye depending on the inhibitor. (M, N, O, P, Q) Cell mortality following rituximab treatment in JOK-1 cells (M), in MEC-1cells (N), in RAMOS cells (O), in two primary B-CLL cells. (P, Q). Cell mortality following venetoclax treatment in RAMOS cells (R), JOK-1 cells (S), in primary B-CLL cells (T). With the exception of primary B-CLL data, all data are representative of at least 3 independent experiments. Statistical significance was analyzed by two-tailed, unpaired Student’s t test.

## 3. Discussion

The role of Orai1 in immune T cell function and other cells such as neutrophils and mast cells and several non-immune cell types was extensively investigated [39]; however, its functional effect in B cells as well as its pathogenic role in malignant B cells is unknown [40]. Here, we investigated the effect of pharmacological blocking of Orai1 constitutive activity on malignant B cell function and signaling. We identified some key findings: (i) Orai1 mediates constitutively active Ca^2+^ entry in various human B-cell cancer lines and primary B-CLL cells and both primary and cell lines display enhanced basal Orai1 channel activity and basal Ca^2+^ signaling; (ii) Inhibition of Orai1 channel decreased cell proliferation in malignant derived B cell lines through cell cycle blockade; (iii) Orai1 blockers induced the loss of mitochondrial membrane potential and, consequently, enhanced B-cell cancer death in caspase-3 independent manner; (iv) Orai1 inhibitors reduced migration of two malignant B-cell lines in dose dependent manner; Orai1 blockers are differentially implicated in B-CLL drug-induced cell death.

Several reports have shown aberrant Ca^2+^ signaling in cancers where mutations in molecules controlling Ca^2+^ homeostasis have been associated with an increased incidence of tumors [41–43], but many key elements of that association are not fully understood. In lymphocytes, Ca^2+^ signaling through CRAC channels is a versatile secondary messenger that regulates cell survival and proliferation by activating or inhibiting a variety of signaling pathways through Ca^2+^ -dependent proteins [28,30]. A similar role of Orai1 in B lymphoblast leukemia, however, has not been reported. Our results herein establish a central role of Orai1 channel activity in malignant B-cells such as B-CLL. Aberrant Ca^2+^ alteration in B-CLL cells is well demonstrated through our work and that of others, which is correlated with patients disease progression [17,26,27,44]. While elevated basal Ca^2+^ levels in B-CLL cells compared to normal B cell has previously been described [27], we show using Orai1 blockers that the constitutive activity of Orai1 is essential for maintaining the basal level of cytosolic Ca^2+^ in B-CLL. Our data provided evidence that Orai1 contributes to SOCE and BCR activities in malignant B-cells such B-cell lymphoma and leukemia lines. This contribution is consistent with a recent study reporting that Orai1 contributed to the majority of SOCE in mouse B cells [13,14,22] but also Ca^2+^ signals driven by antigen-receptor stimulation in mice A20 B cells [14]. Our study also confirms that Orai1 is an essential regulator of B-cell cancer proliferation and viability, in agreement with the study by Gwack **et al.** who observed an inhibition of B cell proliferation from Orai1^-/-^ mice [13] or by Berry **et al.** in naive splenic B cells [28] and by Latour **et al.** who demonstrated toxicity of BTP2, another Orai1 inhibitor, B-CLL samples [45]. However, Emrich and colleagues associated a significant reduction in mouse B cell proliferation and survival with a combined loss of Orai1 and Orai3, whereas this inhibition of proliferation is marginal in B cells from Orai1^fl/fl^ Mb1-Cre/^+^ mice [14]. Our data and the study of Chen et al. [46] suggested that blocking or silencing of Orai1 pathway slowed cell proliferation by controlling the cell cycle G1/S checkpoint. In addition, Orai1 has been shown to be important for cancer cell migration, where its inactivation decreases both cancerous (breast, colorectal, glioma) [34,47,48] or not (VSMCs, HEK293) [49,50] cell migration and metastasis. Supporting these findings, we observed that blocking Orai1 in B-tumor cells reduces their migration capacity. Furthermore, apoptotic resistance effects induced by Orai1 have been described in other cancers. Thus, the role of Orai1 channel in the action of chemotherapy drugs on cancer cells, such as increasing their sensitivity to induce cell death, has been reported e.g. in pancreatic adenocarcinoma cells [51], HepG2 hepatocarcinoma cells [52], acute promyelocytic leukemia cells [53] and ovary carcinoma cells [54]. Consistent with these results, we demonstrated that pre-incubating primary B-CLL cells and B-cancer cell lines with Orai1 blockers increases cell death when associated with drugs. This is also consistent with our previous results showing that blocking constitutive Ca^2+^ entry with antibodies targeting membrane STIM1 increased the sensitivity of rituximab [26].

Overall, the role of Orai1 channel in healthy and pathological B cell function as well as humoral immunity is far from clear, but our study paves the way for further investigation. Indeed, the study of SOCE on malignant B-cells such as B-CLL constitutes an important area of investigation, for example because BCR activation is believed to play an important role in the survival and **in vivo** behavior of B-CLL cells. It can therefore be postulated that by inhibiting this SOCE, antileukemic effects may be obtained, and that Orai1 inhibition may constitute a potentially interesting anti-CLL pharmacological modality, plausibly in combination with conventional drugs used in this pathology, and this is in accordance with data presented in this work. Also it could be the channel that plays a crucial and exclusive role in the functioning of B cell CRAC channels, despite the presence of Orai3, another isoform whose involvement seems to be unnecessary [55]. This study documented a central role of Orai1 in B-cell cancer function such as viability, proliferation, migration and drug sensitivity. Therefore, combination therapy with Orai1 modulators could be used to treat B-cell malignancies in the future.

## 4. Materials and Methods

### 4.1. Cell lines, Cell Culture and reagents

The human B-CLL cell line MEC-1 (DSMZ®, Deutschland), the B-cell prolymphocytic leukemia JVM-3 (DSMZ®), the hairy B cell leukemia line JOK-1 (kindly supplied by Dr. R. Schwartz-Albiez, Heidelberg, Germany [56]) and the Burkitt lymphoma cell line RAMOS (ATCC) were used in this study. MEC-1 cells were cultured in complete IMDM medium, while JVM-3, RAMOS and JOK-1 were cultured in RPMI 1640 medium containing 10% fetal bovine serum, and antibiotics (penicillin and streptomycin) (Sigma®, France), and maintained at 37°C in a 5% CO_2_ humidified atmosphere. Fura-2/QBT was purchased from Molecular Devices, UK. Thapsigargin and Manganese were obtained from Sigma®, France.

### 4.2. Human CLL patient samples

Primary cells from five patients (Table 1) diagnosed with CLL samples (peripheral blood) and three healthy volunteers at the Brest University Hospital were used. Patient consent was obtained and the study was approved by the ethics committee of the Brest University Hospital in accordance with the Declaration of Helsinki. The CLL patients used in this study are part of a prospective registry of CLL patients from the Finistere region, France (https://clinicaltrials.gov/ct2/show/NCT03294980) as previously described [26].

**Table 1.** Clinical characteristics of B-CLL patients included in the study.

| **Characteristics** | **B-CLL patients**<br>**(n=5)** |
| --- | --- |
| Sex | F : 1<br>M : 4 |
| Age at diagnosis | 71y ± 11 |
| Binet classification | A : 3<br>B : 2 |
| IgHV status | Unmutated : 1<br>Mutated : 4 |
| Orai1 expression by flow<br>cytometry<br>(MFI Orai1 - Isotype) | 1.47 ± 0.32 |

Y: year ; IgHV : immunoglobulin heavy chain variable region gene ; MFI : Mean Fluorescence Intensity ; CLL : chronic lymphoblastic leukemia ; F : female; M : male.

Whole blood samples were collected in EDTA-treated tubes and peripheral blood mononuclear cells (PBMC) were isolated by Ficoll-Hypaque density gradient centrifugation (Sigma, France) and B-CLL cells were further purified using the Pan B-cell Isolation Kit (Miltenyi Biotec GmbH, Bergisch Gladbach, Germany). Versalyse buffer (Beckman-Coulter) was added in order to lyse red blood cells for 10⍰min. Only samples where⍰>⍰95% of the cells were CD19⍰+⍰were considered. Cells were maintained in RPMI 1640 medium supplemented as described above, and viability or flow cytometry assays were conducted as described below.

### 4.3. B-CLL drugs and Orai1 inhibitors

Rituximab (anti-CD20), Ibrutinib (Btk inhibitor), Idelalisib (CAL-101) and Venetoclax (ABT-199) were purchased from Selleckchem™ (Selleckchem.com, Houston, TX). Orai1 inhibitors (Synta66 and GSK-7975A) were purchased from Glixx Laboratories, USA. Inhibitors and drugs (except Rituximab) were dissolved and stored in dimethyl sulfoxide (DMSO) at −20°C. Rituximab was diluted and stored in PBS buffer at 4°C.

### 4.4. Orai1 cell surface detection by Flow cytometry

For analysis of cell-surface Orai1 expression, Cells were stained for surface antigens with anti-Human Orai1 (extracellular)-FITC antibody (Alomone labs), according to the manufacturer’s dilution. The determination of the mean fluorescence intensity (MFI) of the Orai1 signal required a minimum of 5000 events. The results were standardized to those obtained with IgG isotype control-FITC. Data were analysed using Kaluza 1.5 software (Beckman-Coulter).

### 4.5. Intracellular Ca^2+^ levels measurements

Cells were labelled with 2⍰μM Fura-2 QBT (Molecular Devices, UK) for 60⍰min at 37⍰°C in buffer containing (in mM, 5 KCl, 135 NaCl, 1 MgCl_2_, 10 HEPES, 1 NA_2_HPO_4_, pH 7.4) and 10 mM glucose and 1.8 mM CaCl_2_, in the presence or absence of Orai1 inhibitors on Cell-Tak (Corning, NY) precoated 96-well plates. Fluorescence acquisition (340 and 380⍰nm excitation wavelengths) and emission (510⍰nm) were analyzed using a FlexStation 3 microplate reader (Molecular Devices). After 100⍰s of recording, cells were stimulated with polyclonal goat anti-human IgM (10 µg/ml) or thapsigargin (2⍰μM) and SOCE was analyzed after readdition of 1.8⍰mM Ca^2+^ (CaCl_2_) at 750⍰s. TG-induced calcium entry was acquired in Ca^2+^-free solution (CaCl_2_ was replaced with 100⍰μM EGTA). SOCE was quantified by the peak of the F_340_/F_380_ ratio. For anti-IgM induced Ca^2+^ response, the extracellular [Ca^2+^]_i_ was 10⍰mM CaCl_2_, before reading, and 10⍰ µg/ml anti-IgM (Jackson Immunoresearch) were injected after 125⍰s. Ca^2+^ entry was quantified after value normalization (ΔF_340_/F_380_).

### 4.6. Basal cytosolic Ca^2+^ measurements

Basal Ca^2+^ levels were monitored with the cytosolic Ca^2+^ indicator Fura-2 QBT. Cells were loaded for 60⍰min with 2⍰µM Fura-2 QBT at 37°C in Ca^2+^ buffer solution (in mM, 5 KCl, 135 NaCl, 1 MgCl_2_, 10 HEPES, 1 NA_2_HPO_4_, pH 7.4) and 10 mM glucose and 1.8 mM CaCl_2_, in the presence or absence of Orai1 inhibitors. Fluorescence was monitored on a FlexStation 3 multi-mode microplate reader by alternately exciting the Ca^2+^ indicator at 340 and 380⍰nm and collecting emitted fluorescence at 510⍰nm. Basal Ca^2+^ levels were estimated as the average of initial F_340_nm/F_380_nm values.

### 4.7. Mn^2+^ quenching

For measurements of Mn^2+^ quenching, cells loaded with Fura-2 QBT were transferred to Ca^2+^-free solution and stimulated with 10 µM Mn^2+^ and fluorescence was measured at an emission wavelength of 360⍰nm before and after Mn^2+^ addition.

### 4.8. Cell viability assay

Cultured B-CLL cells were seeded in culture plates (96 well) with or without Orai1 inhibitors and drugs. After 48 hours, cell viability was assessed by using Live and Dead Cell Vitality Assay (Abcam, UK). Cells were incubated with the LIVE/DEAD kit reagents for 10 min at 37°C and samples were analyzed by flow cytometry with Beckman-Coulter System.

### 4.9. CTV Cell Proliferation assay

CellTrace™ Violet Cell Proliferation (Life Technologies, USA) assay was used to assess cell proliferation by flow cytometry. Cells were labeled with CellTrace™ Violet dye (1:1000 PBS) for 10 minutes, protected from light at room temperature and resuspended in pre-warmed complete culture medium in the presence of Orai1 inhibitors. After 5 days, cells were subjected to flow cytometry with Beckman-Coulter System. We also assessed cell proliferation by using Cell Counting Kit-8 (CCK-8) assay (Sigma, France) according to the instruction, and the absorbance was read at 450 nm under Flexstation-3 Molecular Devices (UK).

### 4.10. Mitochondrial membrane potential (ΔΨ_m_)

The cationic dye JC-10 (a fluorescent ΔΨ_m_ dye, Santa Cruz Inc., Germany) was used to assess mitochondrial membrane potential. JC-10 becomes green when mitochondria are depolarized and red in polarized mitochondria. Cells were incubated with Orai1 inhibitors for 24 hours before loaded with JC-10 (20 μM) for 30 min at 37°C. Then, the medium was substituted with PBS and the fluorescence intensity was measured using flow cytometry with Beckman-Coulter System.

### 4.11. Caspase-3 activity

As reported and described previously [57], activity of caspase-3 was performed by using Ac-DEVD-AFC substrate (Enzo Life Sciences, France). Briefly, B cell lines and primary B-CLL cells were treated with or without Orai1 inhibitors for 24 hours, and then collected and lysed in cell lysis buffer (HEPES 50 mM, NaCl 100 mM, DTT 10 mM, CHAPS 0.1%, EDTA 1 mM, pH 7.4). Cell lysate (20 µg) was added to 100 μL of caspase-3 buffer containing 40 μM of the caspase-3 substrate Ac-DEVD-AFC (fluorogenic) as final concentration and incubated at 37°C for 1 h in the dark. Caspase-3 activity was assessed by measuring fluorescence (Ex 400 nm/Em 505 nm) using Flexstation-3 Molecular Devices (UK).

### 4.12. Apoptosis assay

B-CLL cells were treated for 48 hours in the presence or absence of inhibitors with or without drugs. Apoptotic rate was assessed by annexin V-FITC/PI staining (BioLegend, France). Annexin V-FITC- and propidium iodide-positive cells were determined by flow cytometry using Beckman-Coulter flow cytometer (USA). All flow cytometry data were analyzed using Kaluza 1.5 software.

### 4.13. Cell Cycle Flow Cytometry

After 48 hours of treatment with or without test compounds, the treated B-CLL cells were washed with cold PBS and fixed at room temperature for 30 mn with cold ethanol (70 %). Cells were then washed and incubated for 15 mn in the dark with RNase A (1 mg/ml) and propidium iodide (PI, 1 mg/ml). Cell-cycle distribution was assessed by PI staining using flow cytometry and analyzed by Kaluza 1.5.

### 4.14. Transwell migration assay

B-CLL cell migration rate was determined by using the transwell migration assay (Fisher Scientific S.A.S., France) according to the manufacturer’s instructions. Cells were incubated in 1% fetal bovine serum (FBS) in the presence or absence of Orai1 Inhibitors for 90 minutes. Cells were then placed in the migration upper chambers with or without 30 nM SDF-1 with 1% FBS in the lower chambers. After 24 hours of incubation, migrated cells to the lower chambers were counted as reported elsewhere [58] with fluorescent microbeads (Flow-Count Fluorospheres, Beckman Coulter) by flow cytometry.

### 4.15. Analysis of public datasets

The gene expression dataset used in this study was downloaded from our RNA-seq dataset available under GEO using the following accession number GSE176141 for gene expression analysis of primary samples from Human B-CLL at diagnosis and three years after diagnosis.

### 4.16. Statistical analysis

Statistical analyses were assessed with the PRISM Software (GraphPad Software, (La Jolla, CA, USA) using unpaired t test. Data are expressed as mean⍰±⍰standard error of the mean (SEM) values and p<0.05 were considered statistically significant.

## Author’s contributions

SAA conceived and designed the study, performed the experiments, analyzed and interpreted the data, prepared the figures, and wrote the manuscript. JS performed experiments. CB, BC, YR, and OM reviewed the manuscript. SAA and OM acquired funding. SAA supervised the study.

## Acknowledgments

This work was supported by grants from the Fondation de France (AOC 2017_ NE_ 00078456).

## Data availability

The data used and/or analyzed during the current study are available from the corresponding author on reasonable request.

